# Comparative therapeutic potential of ALX-0171 and palivizumab against RSV clinical isolate infection of well-differentiated primary pediatric bronchial epithelial cell cultures

**DOI:** 10.1101/800326

**Authors:** Lindsay Broadbent, Hong Guo Parke, Lyndsey J. Ferguson, Andrena Miller, Michael D. Shields, Laurent Detalle, Ultan F. Power

## Abstract

Respiratory syncytial virus (RSV) causes severe lower respiratory tract infections in young infants. There are no RSV-specific treatments available. Ablynx has been developing an anti-RSV F-specific Nanobody®, ALX-0171. To characterise the therapeutic potential of ALX-0171 we exploited our well-differentiated primary pediatric bronchial epithelial cell (WD-PBEC)/RSV infection model, which replicates several hallmarks of RSV disease *in vivo.* Using 2 clinical isolates (BT2a; Memphis 37), we compared the therapeutic potential of ALX-0171 with palivizumab, which is currently prescribed for RSV prophylaxis in high-risk infants. ALX-0171 treatment (900 mM) at 24 h post-infection reduced apically released RSV titers to near or below the limit of detection within 24 h for both strains. Progressively lower doses resulted in concomitantly diminished RSV neutralisation. ALX-0171 was approximately 3 fold more potent in this therapeutic RSV/WD-PBEC model than palivizumab (mean IC_50_ = 346.9-363.6 nM and 1048-1090 nM for ALX-0171 and palivizumab, respectively), irrespective of the clinical isolate. When viral genomic copies (GC) were measured by RT-qPCR, the therapeutic effect was considerably less and GCs were only moderately reduced (0.62 – 1.28 Log_10_ copies/mL) by ALX-0171 treatment at 300 and 900 nM. Similar findings were evident for palivizumab. Therefore, ALX-0171 was very potent at neutralising RSV released from apical surfaces but only had a limited impact on virus replication. The data indicate a clear disparity between viable virus neutralisation and GC viral load, the latter of which does not discriminate between viable and neutralised RSV. This study validates the RSV/WD-PBEC model for the pre-clinical evaluation of RSV antivirals.

## Introduction

Respiratory syncytial virus (RSV) is a member of the *Pneumoviridae* family, *Orthopneumovirus* genus (1). It is the leading cause of severe lower respiratory tract infections (LRTI) in infants worldwide (2, 3) with an estimated 33.8 million LRTI cases yearly. RSV accounts for approximately 3.4 million hospitalizations and up to 199,000 deaths worldwide, predominantly in developing countries (4). Economic burden and childhood morbidity and mortality rates associated with RSV are, in many countries, comparable to influenza(5).

Severe RSV infection is associated with an increase in mucus production and a decrease in the number of ciliated cells in the airway epithelium. There is a large influx of immune cells to the airways, predominantly neutrophils but also lymphocytes and macrophages (6). The cellular infiltrate, together with mucus and sloughed epithelial cells, causes lumen obstruction and inflammation of the airways. Associated with mucus plug formation and bronchiole occlusion, bronchiolitis is therefore more severe in smaller airways, such as those of young or preterm infants (7). Accordingly, 66% of RSV-related hospitalizations are in children <6 months old (8). Risk factors associated with the development of severe RSV-LRTI in infants include: prematurity, bronchopulmonary dysplasia, congenital lung or heart conditions, male gender, age ≤6 months, neuromuscular disorders, and immunodeficiency (9). However, the majority of patients that require hospitalization due to severe RSV-related disease have no underlying health conditions that constitute a risk factor (3). There is mounting evidence to suggest that a severe RSV infection in early life is associated with the development of wheeze and subsequently asthma(10).

RSV infection remains a major unmet medical treatment need. Other than the antiviral ribavirin, there is no licensed RSV vaccine or therapeutic, despite the considerable medical importance of this virus. Palivizumab, a neutralizing monoclonal antibody that recognizes a conserved epitope in the viral fusion surface glycoprotein (RSV F site II), is administered prophylactically to high risk infants, e.g., chronic lung disease of prematurity, congenital heart disease or premature birth (typically limited to those less than 29 weeks gestational age for cost/benefit reasons). This is an expensive approach, costing $6,000 to $20,000 per patient for 1 RSV season(11). In addition to price, as indicated above, a major limitation of this approach is that the majority of infants hospitalized with RSV do not fall into these high-risk categories. Palivizumab was assessed as a therapeutic treatment in patients hospitalised with RSV but failed to demonstrate a reduction in viral titers from nasal aspirates or disease severity(12). Therefore, understanding how RSV causes disease in humans and development of therapeutics remains an important medical objective.

One potential limitation to RSV antivirals being effective is that the viral load may have peaked by the time infants are hospitalized. However, a study of RSV clearance in hospitalized children demonstrated that higher viral titers at day 3 of hospitalization were not associated with risk factors such as weight, gestational age, sex, or age at time of admission but were associated with the requirement for intensive care and respiratory failure, indicating a potential therapeutic window even in hospitalized infants (13). Osteltamivir (Tamiflu), an antiviral against influenza virus, demonstrates the importance of the time of administration following infection for effective treatment; it is effective at reducing the length of illness in patients hospitalised with influenza when administered within 48 h of symptom onset in clinically confirmed cases of influenza(14). When administered after this time, however, osteltamivir failed to have any effect on virus titers, disease severity or illness duration(15).

The majority of RSV pathogenesis, antiviral and prophylaxis studies have been performed in animal models or continuous cell lines, neither of which are optimal. Animal models, especially mice, are semi-permissive for RSV replication and do not exhibit high viral titers or pulmonary pathology associated with RSV in infants unless very high innocula are employed (16–18). Continuous cell lines, e.g. HEp-2 and A549 cells, are poorly representative of the complexities of cell interactions in the human lung. The development of the well-differentiated primary pediatric bronchial epithelial cell (WD-PBEC) culture model has provided an authentic surrogate facilitating the elucidation of mechanisms of RSV pathogenesis in pediatric airways (19, 20), and thereby, the study of RSV-specific antivirals. Several studies have demonstrated RSV neutralisation in human airway epithelial cells (HAE) that were not evident in experiments using continuous cell lines such as HeLa or HEp-2(21, 22).

Ablynx has been developing a potential RSV therapeutic, ALX-0171, which is a trivalent non-half-life extended Nanobody® that specifically binds to RSV F site II. ALX-0171 is composed of 3 heavy chain variable region (VHH) domains, providing strong binding to RSV and it potently neutralises infectious virions (23). The neutralizing and therapeutic ability of ALX-0171 against RSV was previously demonstrated in the neonatal lamb RSV-infection model using the RSV strain Memphis 37 (24). In this current study, we exploited our well-differentiated pediatric primary bronchial epithelial cell (WD-PBEC) culture model of RSV infection to assess the relative ALX-0171 IC_50_ for the two RSV clinical strains, BT2a and Memphis 37, in comparison to palivizumab. We also aimed to establish whether any adjustment in the targeted lung lining fluid concentration, as determined in the RSV lamb model, is needed based on strain sensitivity differences.

## Results

There was a clear dose-effect of ALX-0171 treatment on RSV BT2a and Memphis 37 growth kinetics. RSV BT2a titers diminished dramatically by 24 h of treatment with 900 nM ALX-0171 with a 4.74 log_10_ reduction in mean virus titers (Figure 1A). The virus was almost completely neutralized by this time under these conditions and virus neutralization was maintained for the duration of the experiment. Treatment with 300 nM also resulted in reductions in apically-released RSV BT2a, with >2 log_10_ reduction in mean virus titers evident at 144 hpi relative to buffer-treated RSV BT2a-infected control cultures. A less marked reduction in mean RSV BT2a titers was observed following treatment with 100 nM relative to buffer-treated controls.

**Figure 1.**
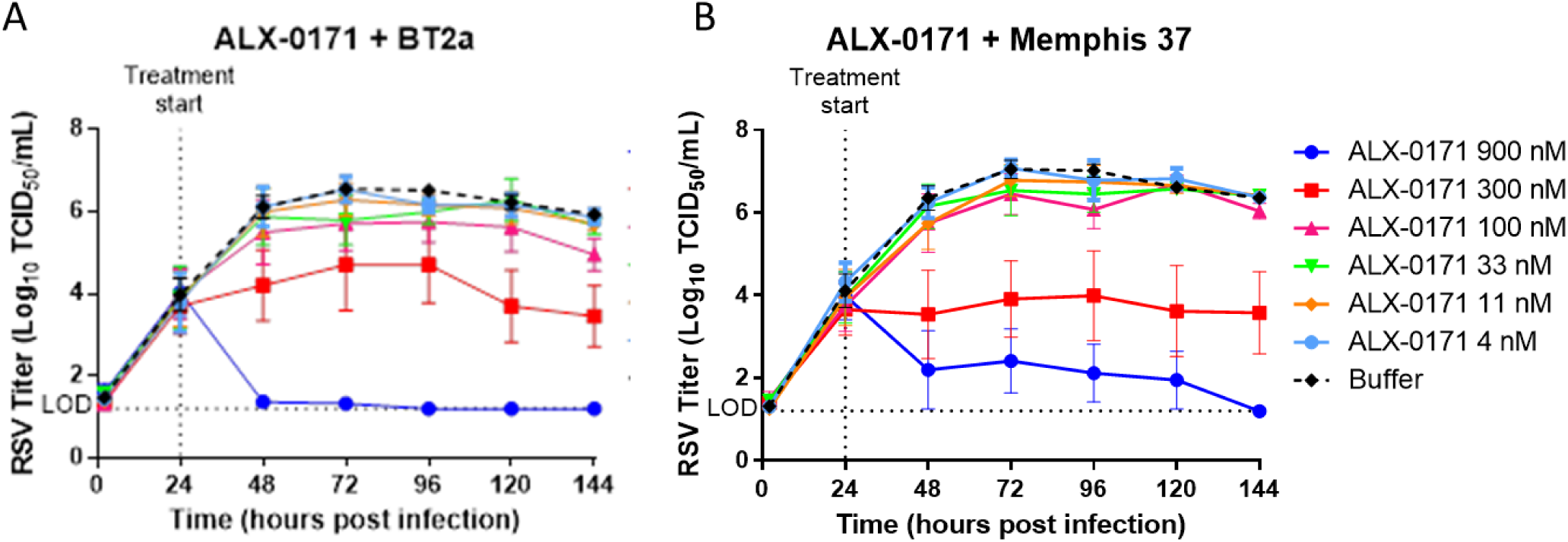
Duplicate WD-PBEC cultures (n=3 donors) were infected apically with either RSV BT2a (A) or RSV Memphis 37 (B) (MOI=0.1) for 2 h at 37°C and then washed 5 times. The fifth wash was retained as the 2 hpi time point for virus titrations. At 24 hpi (and every 24 h thereafter) apical washes were undertaken and harvested for virus titrations. Following the apical washes at 24 hpi (and every 24 h thereafter) the cultures were apically treated with 100 µL ALX-0171 at the indicated concentrations. After 1 h the treatment was removed and replaced with 10 µL ALX-0171 at the same concentration, which remained on the apical surfaces for the duration of the interval between washes. All apical washes were titrated on HEp-2 cells to determine viral titers, which were reported as log_10_ TCID_50_/mL ± SEM (LOD = limit of detection).

Substantial reductions were also evident in RSV Memphis 37 titers by 24 h of treatment with 900 nM ALX-0171, with a 4.2 log_10_ reduction in mean virus titers, although low virus titers were continually detectable in these cultures until 144 hpi (Figure 1B). Lower, but nonetheless substantial, reductions in RSV Memphis 37 titers were also evident following treatment with 300 nM ALX-0171, whereas treatment with 100 nM did not result in substantially reduced viral titers. In contrast, treatment with ALX-0171 concentrations <100 nM did not influence growth kinetics of either RSV strain relative to buffer-treated control cultures under these experimental conditions.

Unlike ALX-0171, infectious RSV BT2a was detected following treatment with all palivizumab concentrations. However, there was a clear dose-effect on virus titers following palivizumab treatment of RSV BT2a-infected WD-PBEC cultures. Treatment with 900 nM palivizumab resulted in >2 log_10_ reduction in viral titers. Treatment with 300 or 100 nM palivizumab also resulted in substantial but lower reductions in viral titers compared to buffer-treated controls (Figure 2A). However, treatment with 33, 11 or 4 nM palivizumab had no effect on the growth kinetics of RSV BT2a.

**Figure 2.**
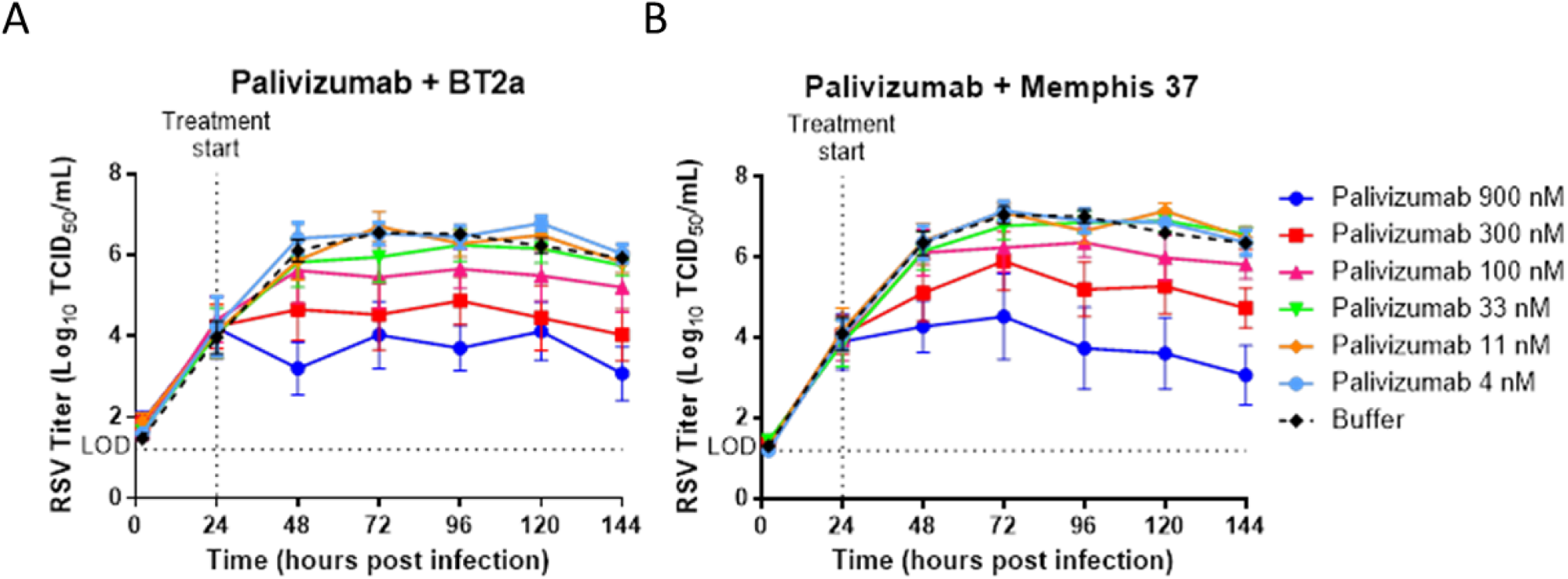
Duplicate WD-PBEC cultures (n=3 donors) were infected apically with RSV BT2a (A) or RSV Memphis 37 (b) (MOI=0.1) for 2 h at 37°C and then washed 5 times. The fifth wash was retained as the 2 hpi time point for virus titrations. At 24 hpi (and every 24 h thereafter) apical washes were undertaken and harvested for virus titrations. Following the apical washes at 24 hpi (and every 24 h thereafter) the cultures were apically treated with 100 µL palivizumab at the indicated concentrations. After 1 h the treatment was removed and replaced with 10 µL palivizumab at the same concentration, which remained on the apical surfaces for the duration of the intervals between washes. All apical washes were titrated on HEp-2 cells to determine viral titers, which were reported as log_10_ TCID_50_/mL ± SEM. (LOD = limit of detection)

Similar results were also observed in RSV Memphis 37 titers following treatment with palivizumab. Treatment with either 900 or 300 nM palivizumab resulted in substantial reductions in viral titers (Figure 2B). However, only slight reductions in mean Memphis 37 titers were evident following treatment with 100 nM palivizumab. Treatment with <100 nM palivizumab did not alter the growth kinetics of RSV Memphis 37 relative to the buffer control.

Treatment with 900 nM, 300 nM or 100 nM of either ALX-0171 or palivizumab resulted in substantial reductions in RSV BT2a titers. At the highest concentration tested (900 nM), ALX-0171 appeared to be considerably more potent than palivizumab, reducing the viral titers to the limit of detection (Figure 3A). The two highest concentrations of both ALX-0171 and palivizumab (900 nM and 300 nM) substantially reduced the viral titers of RSV Memphis 37 compared to buffer-treated controls. However, treatment with ALX-0171 resulted in greater reductions in viral growth kinetics compared to treatment with palivizumab at equimolar concentrations, except for treatment with 100 nM of ALX-0171 or palivizumab, which showed similar viral titers at each time point (Figure 3B).

**Figure 3.**
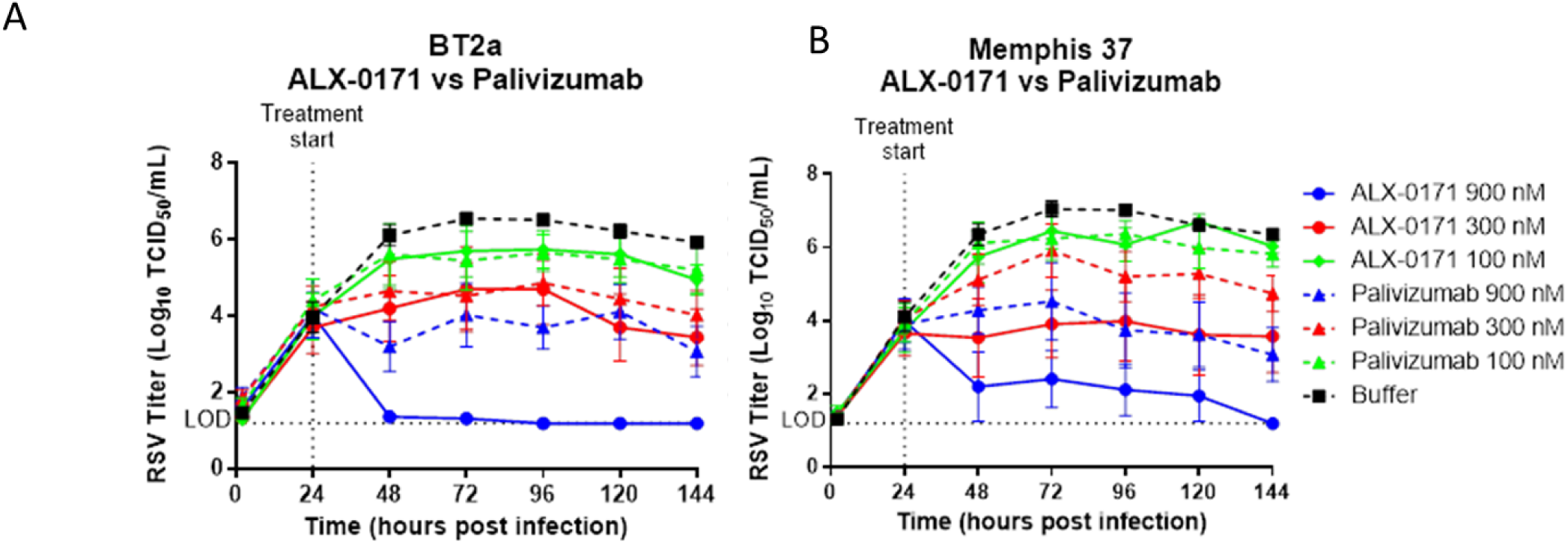
WD-PBEC cultures (n=3 donors and duplicated cultures for each conditions) were infected apically with RSV BT2a (A) or RSV Memphis 37 (B) (MOI=0.1) for 2 h at 37°C and then washed 5 times. The fifth wash was retained as the 2 hpi time point for virus titrations. At 24 hpi (and every 24 h thereafter) apical washes were undertaken and harvested for virus titrations. Following the apical washes at 24 hpi (and every 24 h thereafter) the cultures were apically treated with 100 µL of either ALX-0171 or palivizumab at the indicated concentrations. After 1 h the treatment was removed and replaced with 10 µL of either ALX-0171 or palivizumab at the same concentration, which remained on the apical surfaces for the duration of the intervals between washes. All apical washes were titrated on HEp-2 cells to determine viral titers, which were reported as log_10_ TCID_50_/mL ± SEM. (LOD = limit of detection).

The IC_50_ for each treatment against each virus was calculated (Table 1). The palivizumab IC_50_ (1090 nM) was significantly higher than the IC_50_ of ALX-0171 (363.6 nM) against Memphis 37. Similarly, the IC_50_ of palivizumab (1048 nM) against BT2a was also significantly higher than that of ALX-0171 (346.9 nM). The IC_50_ calculated from the area under the curves was not significantly different for either ALX-0171 ((p-value = 0.751) or palivizumab (p-value = 0.858) against the two RSV strains, indicating that both viruses had comparable sensitivities to neutralisation in the WD-PBEC culture model.

**Table 1:** IC_50_ for ALX-0171 and palivizumab towards RSV BT2a or RSV Memphis 37.

| **RSV Memphis 37** | **IC<sub>50</sub>\* [95% Confidence Intervals] in nM** |
| --- | --- |
| ALX-0171 | 363.6 [276.3-472.9] |
| Palivizumab | 1090 [727.2-2126] |
| P-value (palivizumab v ALX) | 0.0038 |
| **RSV BT2a** | **IC<sub>50</sub>\* [95% Confidence Intervals] in nM** |
| ALX-0171 | 346.9 [262.2-468.3] |
| Palivizumab | 1048 [869-1337] |
| P-value (palivizumab v ALX) | 0.0001 |

A trend towards reduction in RSV BT2a viral loads, as determined by RT-qPCR, was evident following treatment with both ALX-0171 and palivizumab. At 144 hpi, ALX-0171 900 and 300 nM doses reduced mean viral loads by 0.8 log_10_ genome copies (GC)/mL and 1.3 log_10_ GC/mL, respectively, versus buffer–treated cultures (Figure 4A). At the same timepoint and for the same doses, palivizumab reduced mean viral loads by 0.3 and 0.8 log_10_ GC/mL versus buffer-treated cultures, respectively. Similarly, reduced viral loads were observed with both ALX-0171 and palivizumab treatment of Memphis 37-infected WD-PBEC cultures. At 144 hpi, 900 and 300 nM ALX-0171 reduced mean viral loads by 0.62 log_10_ and 0.76 log_10_ GC/mL, respectively, versus the buffer–treated cultures (Figure 4B). At the same time point and for the same doses, palivizumab reduced mean viral loads by 0.27 and 0.57 log_10_ GC/mL, respectively, versus buffer-treated control cultures.

**Figure 4.**
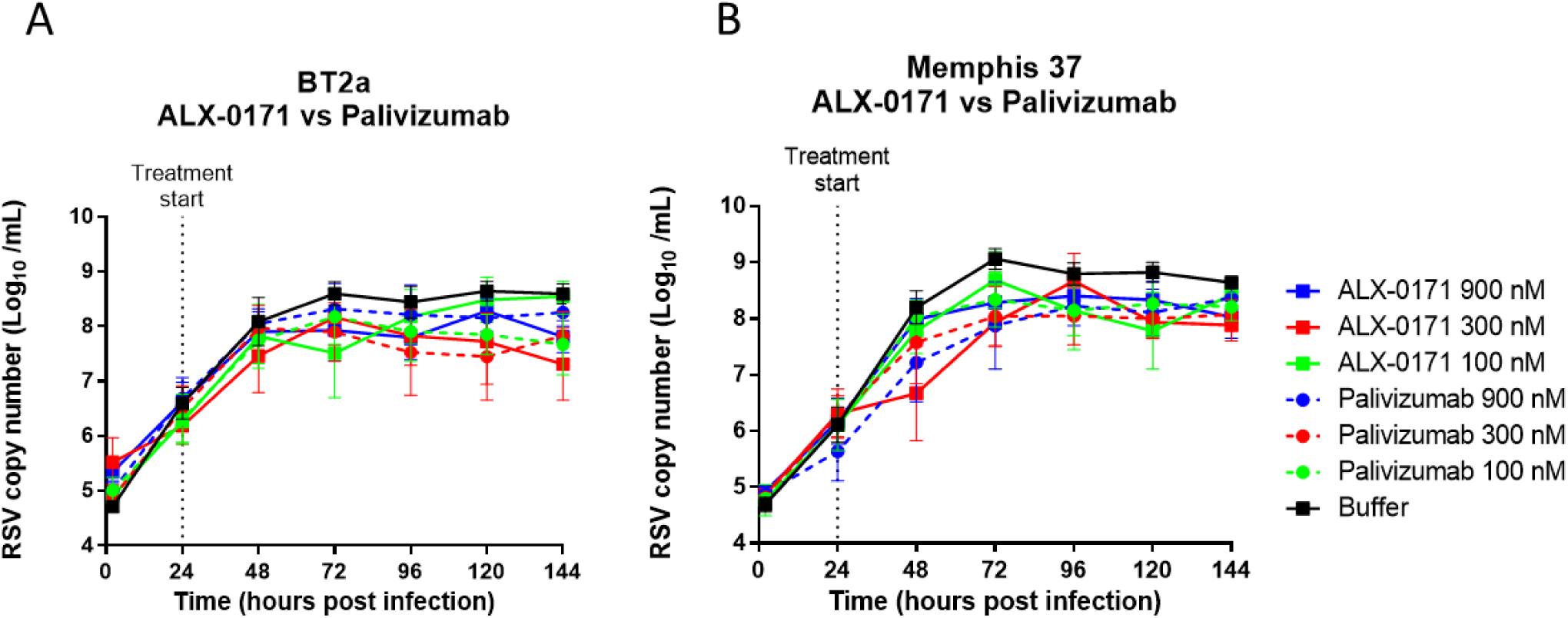
WD-PBEC cultures (n=3 donors and duplicated cultures for each conditions) were infected apically with RSV BT2a (A) or RSV Memphis 37 (B) (MOI=0.1) for 2 h at 37°C and then washed 5 times. The fifth wash was retained as the 2 hpi time point. At 24 hpi (and every 24 h thereafter) apical washes were harvested. Following the apical washes at 24 hpi (and every 24 h thereafter) the cultures were apically treated with 100 µL of ALX-0171 or palivizumab at the indicated concentrations. After 1 h the treatment was removed and replaced with 10 µL of either ALX-0171 or palivizumab at the same concentration, which remained on the apical surfaces for the duration of the intervals between washes. RNA was extracted from apical washes and RT-qPCR performed. Data were plotted as log_10_ genone copies/mL ± SEM.

## Discussion

The aims of this study were two-fold: to assess the efficacy of ALX-0171 as an anti-RSV therapeutic; and to evaluate the WD-PBEC model for use in pre-clinical studies. Despite extensive research into RSV pathogenesis and mechanisms of disease, vaccines and treatments have remained elusive. Ablynx nv has developed ALX-0171 to address the need for a RSV treatment option. In this study we used our RSV/WD-PBEC model to assess the therapeutic potential of ALX-0171 or palivizumab to neutralise two different clinical strains of RSV. Palivizumab is the only licensed neutralising anti-RSV monoclonal antibody. It is administered prophylactically and reserved for infants deemed at high risk of severe RSV infection, in large part because of its high cost. However, the majority of children hospitalised due to severe RSV infection are not classified as high-risk and, as such, do not receive palivizumab prophylaxis. A therapeutic intervention that can be administered after the onset of symptoms would reduce the huge economic burden of RSV and potentially reduce the number and/or duration of hospitalisations.

Growth kinetics for both RSV Memphis 37 and BT2a followed a similar pattern in the WD-PBECs. Both strains reached similar peak viral titers between 72 and 96 hpi. RSV BT2a and Memphis 37 demonstrated similar susceptibilities to neutralisation by ALX-0171 or palivizumab. The highest concentration of ALX-0171 (900 nM) reduced viral titers to near or below the limit of detection by 24 h post-treatment (∼5 log_10_ reduction), whereas palivizumab treatment at the same concentration was less effective (∼3 log_10_ reduction). These differences were reflected in the respective IC_50_ values for both molecules.

The RT-qPCR data demonstrated that ALX-0171 or palivizumab treatment resulted in a trend toward reduced viral replication for both RSV BT2a and Memphis 37 but this did not reach significance. However, the differences were much less marked than the TCID_50_ data. When the TCID_50_ and RT-qPCR data were considered together they suggested that both ALX-0171 and palivizumab treatment resulted in efficient neutralization of RSV released from the WD-PBEC cultures (TCID_50_ results) but had a limited effect on intracellular virus replication. Interestingly, a similar effect was seen following motavizumab administration to infants hospitalised with RSV; a significant reduction in infectious viral titers was reported coincident with a much lower reduction in virus copy numbers(28). However, the RT-qPCR assay does not distinguish between released virus that was neutralized and virus that remained infectious, thereby masking the effect of treatment. Similarly, it was shown that the natural rate of viral load decline is less steep when using a RT-qPCR method than when using quantitative infectivity culture and that this can confound antiviral efficacy determination of test compounds targeting RSV replication(29).

The respiratory system of human infants and young lambs have similarities, suggesting that lambs provide interesting models for asthma, drug delivery, lung development and vaccine efficacy studies(30). A neonatal lamb-RSV Memphis 37 model was also used to assess the efficacy of ALX-0171. There are several similarities in the results from the neonatal lamb model and the WD-PBEC model. Both models showed peak viral titers between 72 to 96 hpi. ALX-0171-treated lambs also demonstrated reduced clinical signs of disease and diminished lung pathology. Importantly, similar IC_50_ values for inhibition of viral growth kinetics following ALX-0171 treatment were obtained from WD-PBECs and neonatal lambs.

The highest Memphis 37 titers reached in the lamb model and the WD-PBEC model in the current study were 4.83 log_10_ FFU/mL(31) and 7.05 log_10_ TCID_50_/mL for the buffer-treated cultures, respectively. As such, RSV evidently reached much higher peak viral titers in the WD-PBEC model compared with the lamb model under these experimental conditions. However, RSV infectivity titers in nasal and/or tracheal aspirates from hospitalised infants were reported to range from ∼10^1^ to ∼10^7^ pfu/mL, suggesting that virus replication in both models may reflect virus growth kinetics in infants(13). In the lamb model viral titers in the lungs were markedly reduced by day 8. However, persistent RSV infection was reported in a WD-PBEC model over a 3 month period, with limited damage to the culture(32). Despite these differences, both the WD-PBEC culture model and neonatal lamb model provided similar IC_50_ values. As the reduction in viral titers was similar in the neonatal lamb model and the WD-PBEC model, and the lambs treated with ALX-0171 had lower clinical severity scores, it is possible that the titer reductions observed in WD-PBECs following ALX-0171 treatment might be predictive of lower clinical severity scores in infants. However, this evidently remains to be confirmed. Nonetheless, our data suggest that our RSV/WD-PBEC model may be of interest in helping to bridge the gap between poorly-predictive pre-clinical animal models and clinical trials to further support the rationale for developing promising RSV therapeutics.

Although palivizumab is licensed for use as a prophylactic, when tested therapeutically it resulted in modest but significant reductions in viral growth kinetics from tracheal aspirates. However, these viral reductions were insufficient to reduce clinical severity in patients hospitalised with RSV(12, 33). Motavizumab, an affinity-matured derivative of palivizumab, which was not approved for prophylactic use, was assessed as a parenterally administered therapeutic following RSV infection. However, there is conflicting data on motavizumab efficacy, with one study indicating a reduction in viral load(28) and another showing no effect on viral load, clinical severity or length of hospitalisation(34). In pre-clinical studies motavizumab and ALX-0171 was 16.8 fold (35) and 126 fold (23)more potent, respectively, than palivizumab. Studies of G-specific antibodies administered post infection have shown a reduction in inflammation in a mouse model of RSV infection(36, 37). This indicates that there is potential for a monoclonal antibody to be used as an effective treatment for RSV, provided the IC_50_ is sufficient. Route of administration is an important consideration. In a study of an adenovirus-based RSV vaccine, intranasal, but not intramuscular, administration elicited strong IgA responses(38). Both palivizumab and motavizumab were administered intramuscularly, whereas ALX-0171 is inhaled. It is likely, therefore that both the increased IC_50_ values and the routes of administration may explain why ALX-0171 appears to have greater therapeutic efficacy *in vivo* than previously developed anti-RSV antibodies.

The determinants of RSV disease severity remain unclear and may involve multiple factors, including viral load, viral strain and host susceptibility. It is also thought that the immune response to RSV infection plays a major role in the severity of disease. High viral titers are associated with higher levels of pro-inflammatory cytokines(13, 39). It has been theorised that a higher viral load may indirectly lead to more severe disease due to an excessive immune response involving production of pro-inflammatory cytokines, leukocyte recruitment and subsequent epithelial cell damage(40, 41). This would correlate with the data from the neonatal lamb model, which showed that reduced clinical severity scores were reflective of reduced viral titers. However, there are other studies that show no correlation between viral load and disease severity(42). It is likely that a combination of host and viral factors contribute to the overall severity of disease.

Interestingly, the neonatal lamb model also resulted in a smaller reduction in virus copy numbers in the airways compared to viral titers following ALX-0171 treatment, despite a significant reduction in clinical severity scores(24). This suggests that the viable virus titer is more indicative of disease severity than the viral load detected by RT-qPCR. Although viral load has been correlated with disease severity in both the human challenge model and infants hospitalised with RSV(43, 44), it may be more appropriate to measure replication competent virus as an indicator of disease severity.

The development of RSV pharmaceuticals presents several challenges. These challenges relate primarily to the fact that much of the pathology associated with RSV infection is thought to be caused by the inflammatory immune responses to the virus infection, rather than direct viral cytopathogenesis. It is imperative, therefore, that RSV antivirals result in both virus neutralisation and modulation of the pro-inflammatory immune responses induced by infection. As has been demonstrated for influenza virus antivirals, such as oseltamivir, early treatment with potent RSV antivirals delivered at sufficiently high doses to the site of infection is likely to be required for effective disease therapy. Using the WD-PBEC model to preform pre-clinical and dose adjustment studies on antiviral treatments offers several benefits. The costs associated with large animal *in vivo* studies is very high and, although WD-PBECs are more expensive than monolayer cell line assays, they can be cultured in a routine class II safety laboratory. Several parameters can be tested in parallel with WD-PBECs and experiments can be carried out in duplicate on cultures from multiple donors. Furthermore, ethical considerations for the use of the neonatal lamb model must include a comprehensive rationale for numbers of animals to be used.

The translation of pre-clinical model data to the clinic has so far proved elusive for RSV therapeutic drugs. The similarity between our RSV/WD-PBEC and the neonatal lamb RSV infection model data suggests that use of our morphologically- and physiologically-authentic RSV/WD-PBEC therapeutic model may provide a basis for predicting drug efficacy in clinical trials that is currently not possible. As such, ALX-0171 may have therapeutic potential against RSV in young infants and our data supports further clinical development.

## Materials and Methods

### Well-differentiated primary pediatric bronchial cell culture

The generation of WD-PBEC cultures was described previously(19). Briefly, primary pediatric epithelial cells obtained from Lonza were expanded in collagen-coated flasks until almost confluent and transferred onto collagen-coated semi-permeable (6 mm diameter, 0.4 µm pore size) Transwells (Corning). When confluent, the apical medium was removed and an air-liquid interface (ALI) was established to promote differentiation. Cells were maintained in ALI for a minimum of 21 days. Cultures were only used when hallmarks of excellent differentiation were evident, including, no holes in the cultures, extensive coverage of beating cilia and obvious mucus production.

### Cell lines and viruses

RSV BT2a was originally isolated from a 4-month old infant hospitalized with bronchiolitis in Belfast, UK. It was cultured in HEp-2 cells as previously described(25) and passaged a total of three times before use. RSV Memphis 37 was originally isolated from a four-month old male in Memphis, USA who presented with bronchiolitis. The virus was isolated and passaged in FDA-approved Vero cell cultures as previously described(26). RSV Memphis 37 was further passaged seven times on HEp-2 cells. RSV titers in biological samples were determined by a tissue culture infectious dose 50 (TCID_50_) assay, as previously described (27), or by RT-qPCR (see below).

### Infection and treatment

WD-PBECs were infected apically in duplicate (2 wells per condition per patient) for 2 h at 37°C. Apical rinses were carried out by adding low glucose DMEM and gently pipetted up and down several times. The recovered DMEM was added to cryovials and snap frozen and stored in liquid nitrogen. Following the 24 hpi apical wash the cultures were treated apically with 100 µL of either ALX-0171 or palivizumab at the indicated concentrations, or buffer control, for 1 h at 37°C. To maintain the air-liquid interface, the treatment was removed and replaced with 10 µL of the same concentration of ALX-0171 or palivizumab and incubated for 24 h, until the next apical rinse. After each subsequent apical rinse 10 µL of the indicated concentration of ALX-0171 or palivizumab were added to the apical surface. This was repeated every 24 h for 6 days.

### Virus Quantification

RSV titers in apical washes were determined on HEp-2 cells as previously described(27). To determine the viral load by RT-qPCR RNA was extracted from apical washes (High Pure Viral RNA kit, Roche). cDNA was prepared using 10 µL RNA (High Capacity cDNA Reverse Transcription kit, ABI). The Light Cycler 480 probe master kit (Roche) was used to amplify RSV cDNA. Primers and probes specific for RSV L-gene were designed with Mega6 software based on alignment of multiple RSV L-gene sequences derived from GenBank representing both clinical and prototypic strains of RSV A and B subgroups (Table 2). Standard curves were generated using a plasmid containing the RSV-A2 genome in 10-fold dilutions.

**Table 2:**
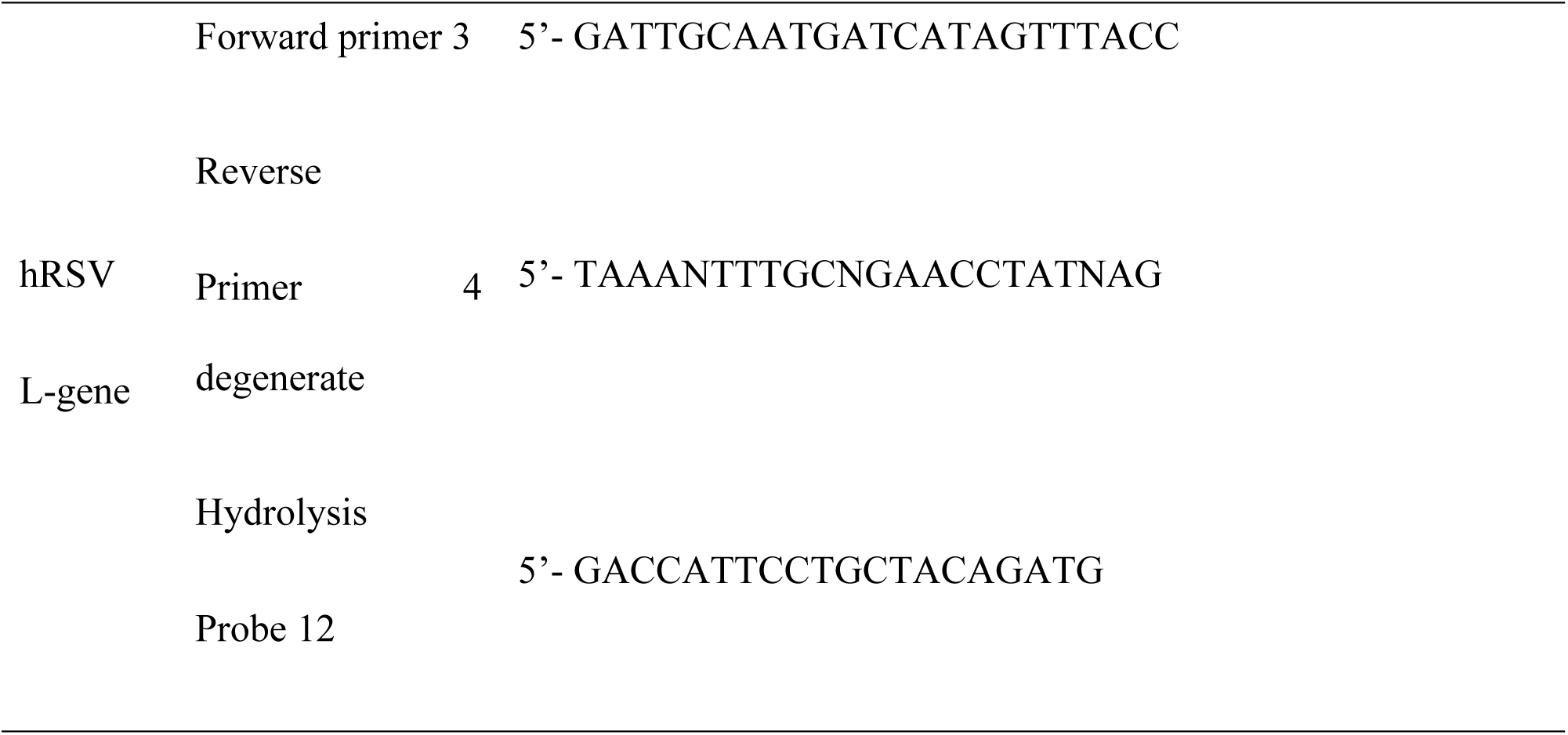
RT-qPCR primer and probe sequences.

### Statistical Analysis

At each time point, 4 parameter logistic (4PL) curves were fitted to the TCID_50_ data for each RSV strain and compound in GraphPad Prism version 7.03. The bottom was constrained to be log_10_ 1.2, i.e. the limit of detection for TCID_50_, and the top was constrained to the mean response of buffer-treated cultures. In order to detect any difference between either ALX-0171 and palivizumab or RSV strains, t-tests were performed based on the IC_50_ from the 4PL model. Since IC_50_ is a lognormally distributed variable, t-tests were performed on the log-scale. For this purpose, the 95% asymptotic confidence intervals were log_10_-transformed and the standard error was derived as the difference between the upper and lower limit divided by 2 times the t-quantile. The change in TCID_50_ values over time was summarized per donor using the area under the curve (AUC) for each compound and concentration. The lower limit of detection of TCID_50_ was used as a baseline for calculation. Subsequently, the dose response of average AUC values was fitted using a 4PL with the bottom constrained to 0. For RT-qPCR data, area under the viral load curves (AUC) were computed using the trapezoid rule in Graphpad Prism for each concentration of either ALX-0171 or palivizumab and compared to the AUC of the buffer-treated cultures using an ANOVA. A post-hoc Dunnett’s test to correct for multiple testing was performed. A p-value <0.05 was considered statistically significant.

## Acknowledgements

This study was wholly funded by Alynx nv, Belgium, in a research project conducted at Queen’s University Belfast.

